# Tailoring hydrophobicity and strength in spider silk-inspired coatings via thermal treatments

**DOI:** 10.1101/2024.08.12.607662

**Authors:** Anni Seisto, Anna S. Borisova, Robert Pylkkänen, Pezhman Mohammadi

## Abstract

The advent of advanced coatings has transformed material functionalities, extending their roles from basic coverage and visual appeal to include unique properties such as self-healing, superior hydrophobicity, and antimicrobial action. However, the traditional dependency on petrochemical-derived materials for these coatings raises environmental concerns. This study proposes the use of renewable and alternative materials for coating development. We present the use of bioengineered spider silk-inspired protein (SSIP), produced through recombinant technology, as a viable, eco-friendly alternative due to their ease of processing under ambient pressure and the utilization of water as a solvent, alongside their exceptional physicochemical properties. Our research investigates the effects of different thermal treatments and protein concentrations on the mechanical strength and surface water repellency of coatings on silica bases. Our findings reveal a direct correlation between the temperature of heat treatment and the enhancements in surface hydrophobicity and mechanical strength, where elevated temperatures facilitate increased resistance to water and improved mechanical integrity. Consequently, we advocate SSIPs present a promising, sustainable choice for advanced coatings, providing a pathway to fine-tune coating recipes for better mechanical and hydrophobic properties with a reduced ecological footprint, finding potential uses in various fields such as electronics.

## 1. Introduction

Advanced material coatings are at the forefront, driving transformative breakthroughs in materials science and engineering. The quest for advanced material functionalities has steered the scientific community towards the exploration of innovative coatings that not only serve protective and aesthetic purposes, but also include materials with properties like self-healing, enhanced hydrophobicity, and antimicrobial capabilities. ^[1–3]^ Central to this pursuit is the shift from traditional petrochemical-based materials towards integrating innovative materials science and sustainable practices. By harnessing the potential of renewable resources and minimizing the dependency on petrochemical derivatives, we can pave the way for a circular economy that prioritizes environmental sustainability without compromising technological advancement.

The utilization of biopolymers in coatings such as proteins, polylactic acid (PLA), polyhydroxyalkanoates (PHA), cellulose nanofibers (CNF), cellulose nanocrystals (CNC), lignin, chitin, chitosan, and alginate is increasingly significant due to their environmental and functional merits. ^[4,5]^ These biopolymers are explored as sustainable alternatives to synthetic polymers because of their biodegradability and reduced ecological footprint. Notably, CNF and CNC derived from cellulose offer exceptional strength and rigidity, enhancing the structural integrity of biocomposite materials coatings.^[6,7]^ Lignin, an abundant natural polymer, contributes to UV resistance and antioxidant properties, broadening the application spectrum to include outdoor and protective coatings.^[8–10]^ Similarly, chitin and chitosan, extracted from shellfish exoskeletons, are lauded for their biocompatibility and antimicrobial properties, making them invaluable in medical and food packaging applications.^[11–13]^ Alginate, sourced from seaweed, excels in gel-forming capabilities, which are crucial for encapsulation and barrier applications.^[14,15]^ The integration of these biopolymers with PLA and PHA can lead to composites with tailored properties such as enhanced durability, stability, and functionality, meeting specific needs across various sectors.^[16–18]^

The versatility of biopolymer coatings is underscored by their ability to create innovative material solutions that are not only environmentally sustainable but also highly effective in diverse industrial and technological contexts. One notable example is the adoption of high-performance bioengineered materials. ^[19–25]^ An alternative approach involves the bioengineering of biopolymer building blocks directly within microbial systems through industrial biotechnology.^[21]^ This method harnesses the capabilities of engineered microbes to synthesize biopolymers to specific functional properties. The advantages of microbial synthesis include the ability to scale production sustainably, reduce dependency on side streams such as agricultural resources, and enhance the uniformity and purity of the biopolymers produced.^[21]^ Moreover, this strategy enables precise control over the molecular structure of biopolymers, which can lead to materials with improved performance characteristics tailored to meet the rigorous demands of various applications, thereby fostering innovation in material science.^[21]^

In the pursuit of next-generation high-performance materials, we propose a shift towards the integration of bio-based components, such as spider silk-inspired proteins (SSIPs). Utilizing recombinant DNA technology, this study showcases a significant stride in this direction by introducing SSIP as a sustainable, high-performance option for advanced coatings.^[21]^ The use of SSIPs for advanced coating applications is motivated by their remarkable self-assembly capabilities and intrinsic material properties, which are leveraged to meet the demands of next-generation materials. SSIPs, exhibit superior mechanical strength, elasticity, and adhesiveness, making them highly suitable for biomimetic applications.^[22,24–27]^ The transition towards utilizing SSIPs in coating technologies specifically addresses the need for sustainable yet high-performance materials in industries like aerospace, automotive, and consumer electronics, where traditional coatings may fail to meet increasingly stringent environmental and functional standards. The integration of SSIPs into thin film technologies enhances this proposition by enabling uniform film deposition, minimal material waste, and reduced environmental impact. Such films consistently demonstrate enhanced properties such as improved barrier functionality, mechanical resilience, and binding stability.

In this work we investigate the interplay between thermal treatments and SSIP concentrations on the coatings’ hydrophobicity and mechanical properties when applied to silica substrates. Through systematic analysis, we discern a complex relationship where both the concentration of SSIPs and the applied thermal treatments serve as critical levers in modulating the surface properties of the resulting coatings. At lower concentrations, an increase in SSIPs leads to a notable enhancement in surface hydrophobicity, evidenced by rising contact angles. This trend reaches an apex at intermediate concentrations, beyond which further increases in concentration paradoxically diminish hydrophobicity. Thermal treatments further compound this complexity, with elevated temperatures generally bolstering hydrophobic properties, particularly at lower SSIP concentrations.

Parallel to the investigation into hydrophobicity, this study extends its analysis to the nanomechanical properties of SSIP coatings, uncovering a similar dependence on concentration and thermal conditions. At optimal conditions, identified through hydrophobicity studies, the coatings exhibit significant enhancements in both modulus and hardness, suggesting a thermally induced densification and possibly increased cross-linking within the protein structure.^[28–30]^ Intriguingly, the study also reveals the heterogenity within SSIP coatings, manifested through variations in color that correlate with film thickness and, consequently, mechanical properties. This aspect introduces an additional layer of complexity, indicating that film thickness—and by extension, the physical structure of the coating—plays a pivotal role in determining the material’s mechanical behavior.

Collectively, these insights illuminate the nuanced dynamics governing the properties of SSIP-based coatings, highlighting the interplay between material concentration, thermal treatment, and physical structure. This understanding paves the way for tailored fabrication processes, enabling the optimization of SSIP coatings for diverse applications that demand specific hydrophobic and mechanical characteristics.

## 2. Material and Methods

### 2.1 Molecular cloning, protein expression and purification

All procedures related to cloning, expression, and purification followed methodologies outlined in previous literature.^[24–27,31]^ The DNA sequences for the bacterial family III cellulose-binding module (CBM3) from *Ruminiclostridium thermocellum*, along with twelve repeated sequences from *Araneus diadematus* major ampullate spidroin 3, and the terminal linkers of major ampullate spidroin 1 from *Euprosthenops australis*, were synthesized with codon optimization for E. coli expression by GeneArt (ThermoFisher Scientific). The assembly of seamless fusion constructs utilized Golden Gate cloning, employing the pE-28a (+) (kanR) vector from Novagen, which includes a C-terminal 6x Histidine-tag, to create the construct named CBM-eADF3-CBM. *E. coli* strain 10-β competent cells, specifically chosen for their genetic makeup conducive to cloning, were obtained from New England Biolabs. The expression utilized either the BL21 strain or the BL21 T7 express™ strain from ThermoFisher Scientific, with standard growth media and conditions unless noted otherwise. Following expression, colonies were initially cultured in LB media with kanamycin at 37°C-250 rpm, then scaled up in oxygen-rich LB media until reaching mid-log phase. Induction was initiated with isopropyl β-D-1-thiogalactopyranoside (Sigma-Aldrich), followed by a temperature reduction to 20°C for extended expression. Cells were subsequently collected by centrifugation.Purification began with resuspending cell pellets in Lysis Buffer, containing various components for cell disruption and protein stabilization, and proceeded with sonication for cell lysis. Post-sonication, centrifugation separated the soluble proteins, which were then purified using HisTrap FF columns on an ÄKTA-Pure FPLC system, following the manufacturer’s guidelines for binding and elution. Desalting was performed using Econo-Pac10 DG columns from Bio-Rad, and proteins were analyzed via SDS-PAGE, stained with Coomassie Brilliant Blue for visualization. Protein concentrations were determined using a DS-11 FX Spectrophotometer at 280 nm absorbance. Finally, for storage, proteins were lyophilized using an Alpha 2–4 LSCBasic lyophilizer (Christ).

### 2.2 Coating formulations

Coating formulations were prepared by dissolving the lyophilized protein powder in MilliQ water to the final concentration of 0.01 - 10 g/L and then centrifuged for 10 min at 10000 rpm (Eppendorf Centrifuge 5430 R). The supernatant was taken for surface coating.

### 2.3 Spin-coating of thin films

Prior to spin-coating, silica wafers (1 × 1 cm^2^; Compugraphic, Jena, Germany) were placed in a UV-ozonizer (UV/OzoneProCleaner, Bioforce Nanosciences, Ames, IA, USA) for 15 min and then wetted with pure toluene, followed by spin-coating the toluene solution. After that 100 µl of SSIP (0.01 - 10 g/L) was added on the top of silica and let stand at RT for 10 min prior spinning. The spin-coating was performed in a Autolab spincoater device (Ekochemie, Germany) at 2000 rpm for 2 min. Spin-coated films were treated at different temperatures (23 - 170 °C) for 10 min in the oven.

### 2.4 Contact angle measurement of wettability

The wettability of the thin films was determined by static water contact angle (WCA) measurements using the sessile drop method in a CAM 200 (KSV Instruments, Finland) goniometer equipped with a video camera at RT. The WCA was measured from the drop shape using the Young−Laplace equation. The WCA was recorded (3 times per second) within 60 s immediately after placing the droplet (2 μL) on the surface of the spin-coated film; the WCA was calculated as the average value of the measurements.

### 2.5 Structural changes

Raman spectra were recorded using an inVia Confocal Raman Microscope (Renishaw, UK) utilizing a 785 nm laser with nominal power of 100 mW at 100% intensity, accumulation from 5 scans (10s), a Leica 0.4 NA 20X objective, and diffraction grating of 830 mm^−1^. The samples were prepared by casting 100 μL of SSIP solution (10 g/l) on aluminum foil and drying at 23 °C (untreated sample) and 110 °C (heat-treated sample).

### 2.6 Nanomechanical measurement

Nanomechanical tests were carried out using an iNano® nanoindenter (KLA Corp., USA) with a Berkovich diamond tip (Synton MDP, Switzerland) equipped with an InForce 50 mN electromagnetic force actuator 100kHz data acquisition rate and 20µs time constant, a dynamic indentation module (continuous stiffness measurement (CSM), a NanoBlitz for high-speed nanoindentation to creating 2D large area mapping module, and a scratch test module. Oliver and Pharr method was used to calibrate the diamond area function (DAF) of the tip as described previously.^[22,32,33]^ For the CSM measurements target load of 50mN, indentation strain rate 0.200 S^-1^, frequency 110 Hz and target displacement of 2 nm was used. For the 2D mapping area of 100 × 100 µm was selected. For the scratch test, we performed the measurement using the scratch length of 150 µm and with the scratch starting load of 0.04mN and maximum load of 20 mN using 20 µm/s scratch velocity.

## 3. Results and discussion

### 3.1 Starting materials and coating fabrication

We examined material behavior of SSIP across a range of concentrations and response to temperature leading to thirty variations. **Figure 1** showcasing a spectrum of coatings formulation prepared from SSIP solution concentrations ranging from 0.01 mg/ml to 10 mg/ml. These coatings were systematically tested across a temperature range from 23°C to a high of 170°C. SSIP solutions were prepared in MQ-water at various concentrations and subjected to vortex and centrifugation to achieve uniformity, omitting any solubilizing agent to preserve the material’s intrinsic properties. The spin-coating technique was then employed to apply SSIP evenly across silicon substrate surfaces, but also prevent the formation of air bubbles and to ensure the integrity of the film structure. Once applied, the coatings were left to dry, forming films that displayed a range of surface characteristics influenced by temperature.

**Figure 1.**
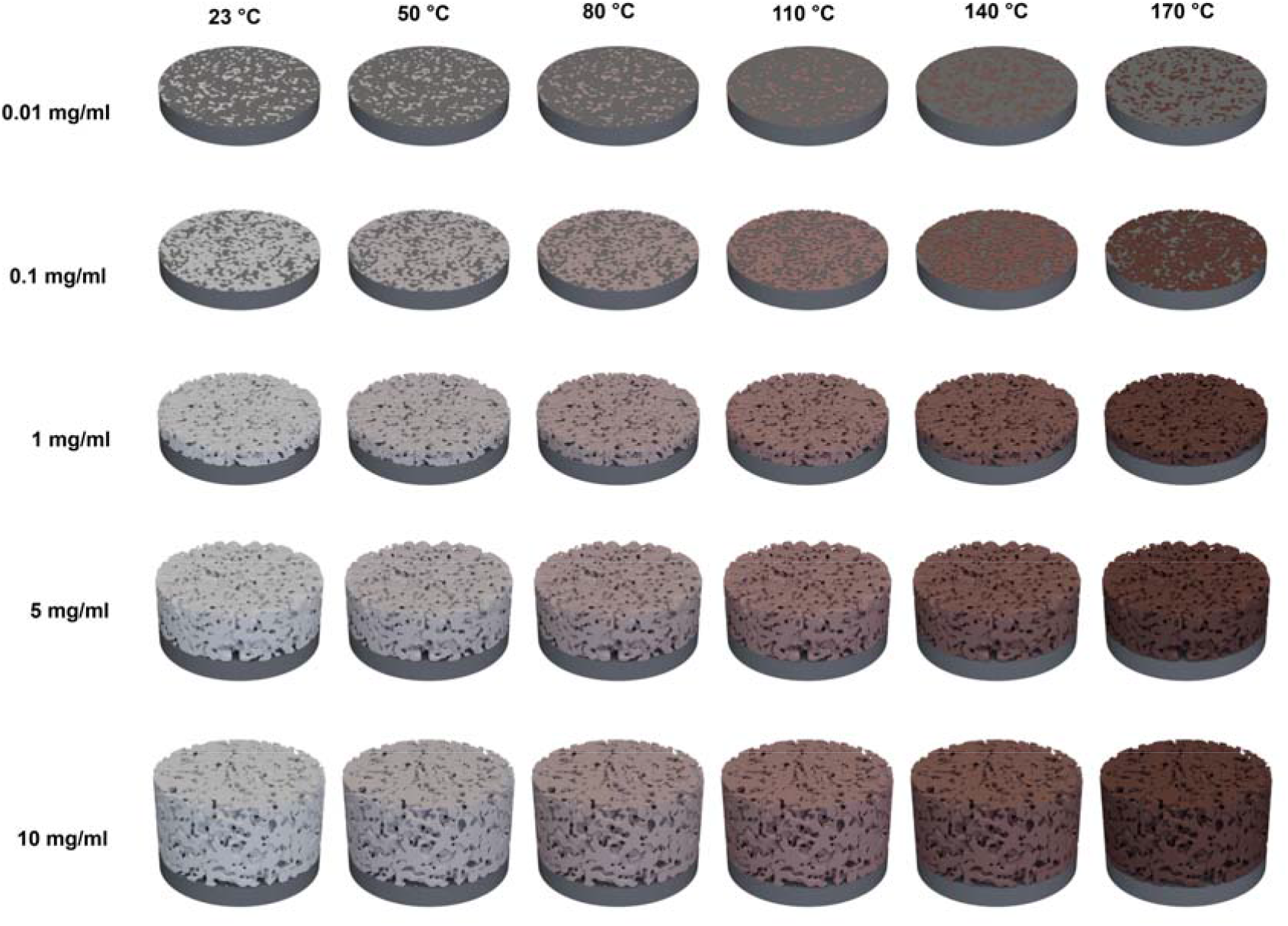
**Schematic representation of SSIP coatings illustrating various concentrations and heat treatments used in this study**.

### 3.2 Thermal modulation of SSIP surface hydrophobicity at varying concentrations

The contact angle measurements for the SSIP across varying concentrations and heat treatment temperatures revealed notable trends and dependencies (**Fig. 2**). At concentrations ascending from 0.01 to 1 mg/mL, there is a trend of increasing contact angles, suggesting an increase in surface hydrophobicity. This increase is pronounced, reaching a peak contact angle that is approximately 50-60% higher than at the lowest measured concentration. Upon surpassing this concentration threshold, the trend inversely correlates, with contact angles decline, indicative of reduced hydrophobicity—exhibiting a decrease to nearly half of the peak values at the highest concentration (10 mg/mL). Looking at the effect of temperature, it becomes apparent that higher heat treatments enhance hydrophobicity, particularly in the lower concentration range. For example, at 1 mg/mL, the contact angle under the 140 °C treatment is nearly double (approximately a 100% increase) that observed at room temperature, referred to in our work as “optimal coating”.

**Figure 2.**
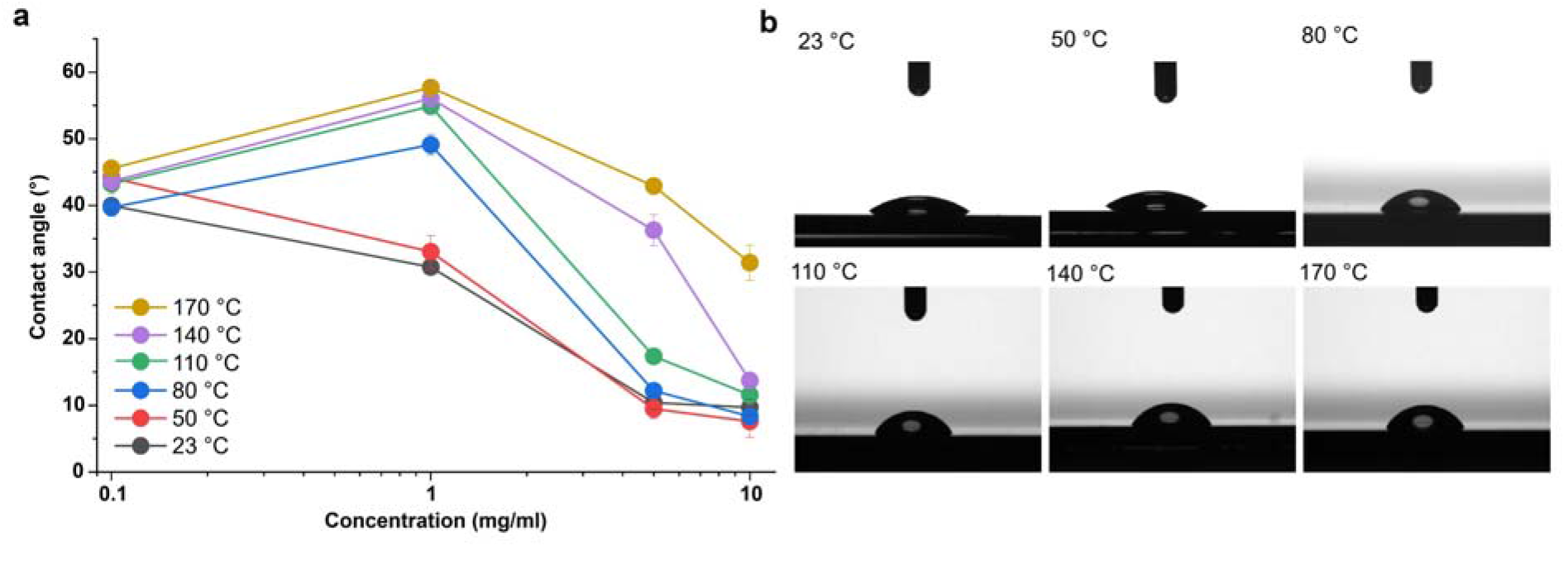
Temperature and concentration-dependent hydrophobicity of the SSIP. (a) Contact angle measurements for SSIP coatings ranging from 0.1 to 10 mg/ml, with temperature increases from 23°C to 170°C. (b) Droplet profiles on 1 mg/ml SSIP-coated surfaces at selected temperatures, illustrating the shift from hydrophilic to hydrophobic behavior with rising temperatures.

Conversely, this trend inverts at higher SSIP concentrations approaching up to 10 mg/mL, where the temperature’s impact diminishes and the contact angles converge, regardless of the heat treatment (**Fig. 2**). This includes RT and 50 °C up to 170 °C, converging to similar contact angle measures, which are significantly lower than the optimal coating—a convergence that translates to a more than 80% reduction from the peak contact angle observed at 140 °C and 1 mg/mL concentration. This convergence at high concentration levels suggests a threshold above which increasing the concentration or altering the heat treatment temperature provides negligible improvement to hydrophobicity. These insights underline the delicate interplay between SSIP concentration and thermal treatment in dictating surface wettability and provide quantifiable benchmarks for optimizing SSIP-based coatings to enhance their hydrophobic properties in a sustainable and controlled manner.

Our investigation’s outcomes strongly suggest that the hydrophobic characteristics of the SSIP-coated surfaces can be modulated by both the protein solution’s concentration and the ambient temperature during the heat treatment process. For the optimal coating, it is conceivable that molecular reorientation effects, as prompted by temperature increments, may lead to a favored redistribution of hydrophobic side chains toward the coating-air interface, potentially enhancing the surface’s hydrophobicity. The observed increase in contact angles, especially at optimal coating, hints at an ideal repositioning of hydrophobic moieties in proximity to the air. One could also speculate that the process of heat-induced protein denaturation, followed by aggregation, could expose more hydrophobic domains, as indicated by the marked upsurge in contact angle measurements, which nearly doubled at 140 °C in comparison to room temperature at a concentration of 1 mg/mL (**Fig. 2**). This phenomenon could suggest a shift toward a structure inherently more conducive to hydrophobicity. Also, the interplay between solvent evaporation and SSIP concentrations, particularly at heightened temperatures, appears to be a variable affecting the density at which protein molecules arrange themselves on the substrate. This could, in turn, impact the hydrophobic attributes. Also, rearrangement and the formation of crystalline structure during heat application might imply a more systematic arrangement of hydrophobic residues, though this remains a matter for further inquiry.

On the other hand, the observed decline in hydrophobicity at elevated SSIP concentrations might be explained by a confluence of factors that require further investigation. For instance, protein overcrowding on the surface could lead to an orientation that inadvertently exposes more hydrophilic residues, negating the hydrophobic interactions essential for higher contact angles. Additionally, the increased thickness and potential porosity of the coating at higher concentrations might trap water molecules, diminishing hydrophobicity. This effect could be compounded by retained solvent within the coating, a consequence of the reduced solvent evaporation efficiency due to the higher solution viscosity at greater protein concentrations. Furthermore, excessive protein might promote non-uniform aggregation or phase separation, introducing hydrophilic patches on the surface and thus lowering the overall hydrophobicity. Conformational changes induced by intermolecular interactions at high concentrations could also expose polar groups to the surface, further reducing hydrophobicity. Lastly, a competitive adsorption phenomenon, where hydrophilic components of the solution might preferentially adsorb to the surface at higher concentrations, could also contribute to this decline. This complex synergistic interplay suggests a delicate equilibrium between factors enhancing and diminishing hydrophobicity, highlighting the various nature of protein-surface interactions at different concentrations.

### 3.3 Influence of heat-treatment on protein structures

To better understand the potential impact of heat treatment on the structural changes and the integrity of the coatings, we carried out Raman spectroscopy, particularly focusing on the amide I, II, and III bands which are indicative of specific protein configurations. Our results, depicted in **Figure 3**, demonstrate distinct changes in the spectral profiles between heat-treated and untreated samples, quantitatively analyzed through Gaussian fitting.^[34]^ The most critical observations were made in the Amide I region, where the peak position for both untreated and heat-treated coating was centered around 1667 cm^-1^ and 1668 cm^-1^, respectively. Notably, the full width at half maximum (FWHM) decreased from 49.6 cm-1 in untreated samples to 58.4 cm^-1^ in heat-treated case.^[34]^ Although there was no significant shift towards the β-sheet spectral range (1630-1640 cm^-1^), the observed reduction in FWHM suggests a denser packing and potentially an increase in β-sheet content, significant since Amide I is sensitive to C=O stretching vibrations associated with these structures. In the Amide III region, the heat-treated silk exhibited a sharper and slightly shifted peak at 1257 cm^-1^ with a FWHM of 85.4 cm^-1^, compared to 1256 cm^-1^ with a FWHM of 81.7 cm^-1^ in untreated sample. This reduction in FWHM indicates a tighter, more ordered arrangement of protein structures, likely reflecting an increase in alpha-helix or β-turn content, which often accompanies β-sheet structures in thermally influenced proteins. Lastly, in the Amide II band, we observed a notable narrowing from 48.4 cm^-1^ FWHM in untreated silk to 42.2 cm^-1^ FWHM in the heat-treated samples, alongside a peak shift from 1450 cm^-1^ in untreated to 1450 cm^-1^ in treated samples. This suggests a more consistent and aligned protein environment, indicative of stronger N-H bending and C-N stretching interactions post-heat treatment. These results collectively suggest that heat treatment may lead to a more ordered and potentially more stable protein structure in SSIPs.

**Figure 3.**
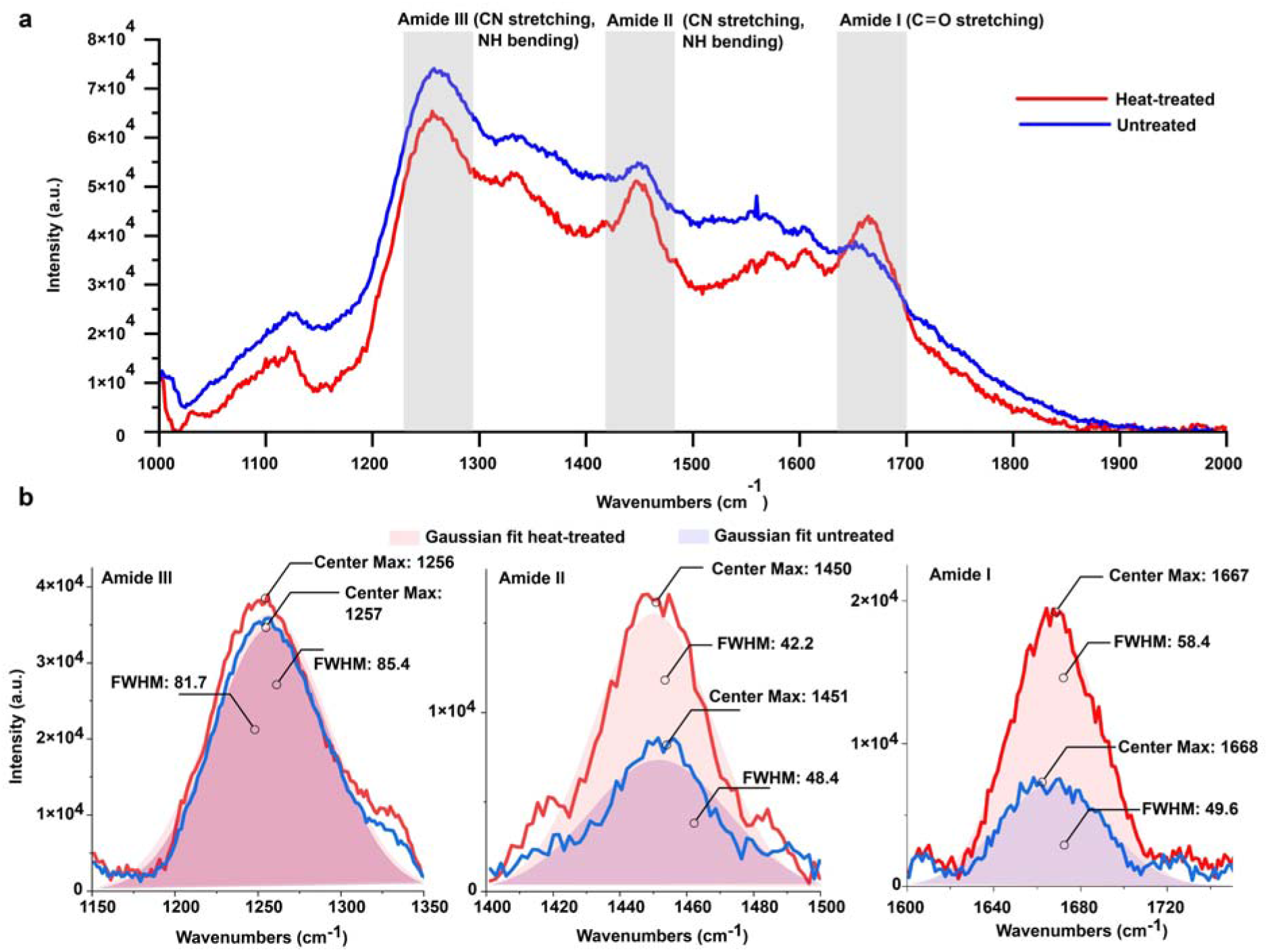
Raman spectroscopy of heat-treated and untreated coating. **(a)** Comparative Raman spectra of the coating with (blue) and without (red) heat treatment, spanning the range of 1000-2000 cm^-1^. The spectra highlight the amide regions: Amide III (1200-1300 cm^-1^), Amide II (1450-1600 cm^-1^), and Amide I (1600-1700 cm^-1^), which are key indicators of protein secondary structure. (b) Gaussian fits for the amide peaks of both heat-treated (blue) and untreated (red) coating. The fits elucidate the changes in peak positions and widths within each amide region.

### 3.4 Thermal modulation of SSIP nanomechanics

In alignment with the previously observed trends in surface hydrophobicity of SSIP coatings, our investigation into the thermal modulation of the nanomechanical properties uncovers analogous dependencies on both concentration and thermal treatment. At the general level examination of SSIP films, following heat treatments ranging from 23 °C to 170 °C, unveils that mechanical modulation, as characterized by modulus and hardness, is significantly influenced by the interrelated factors of temperature and SSIP concentration (**Fig. 4**).

**Figure 4.**
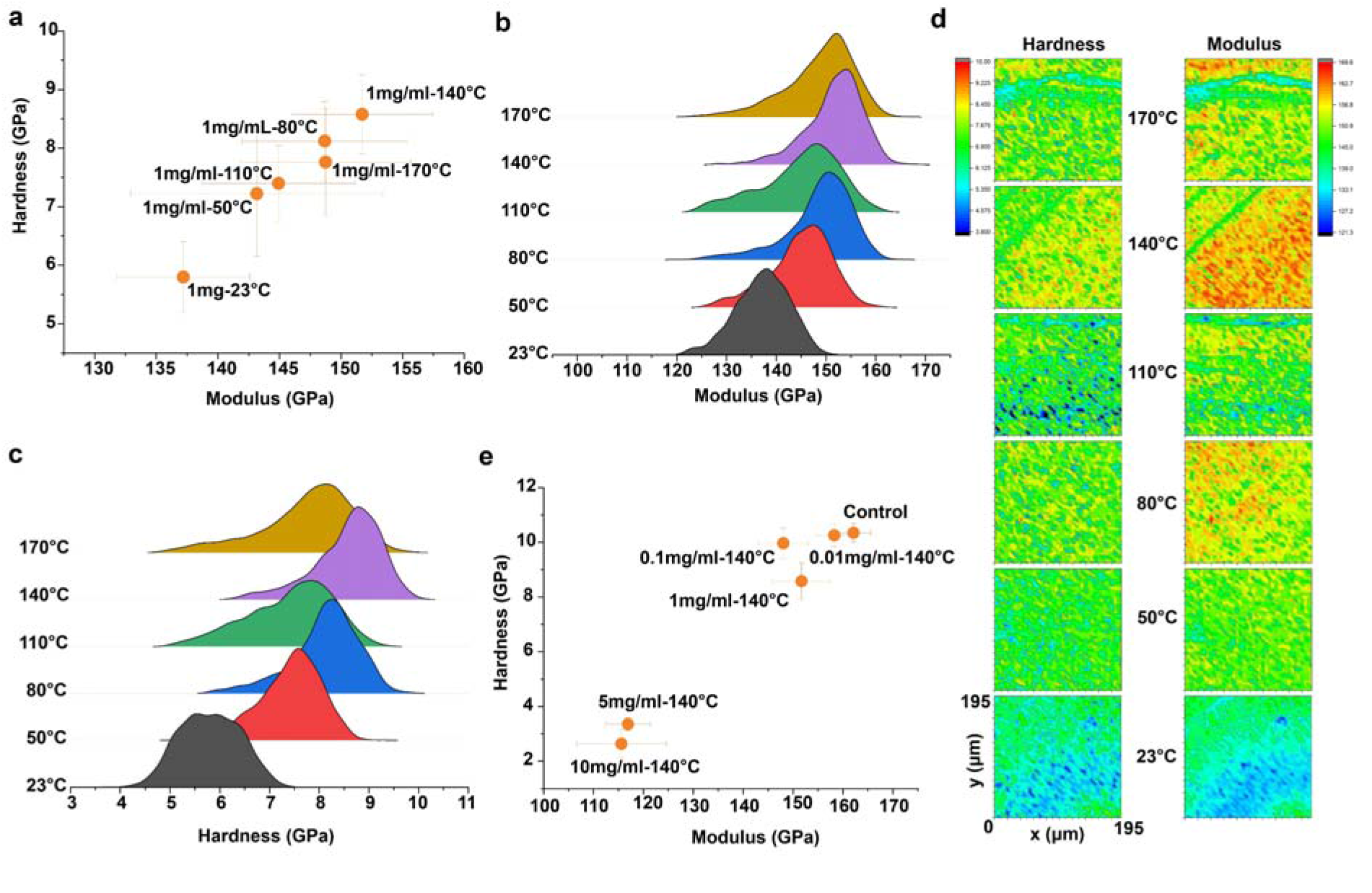
Mechanical properties and surface morphology of SSIP coatings at various temperatures and concentrations. (a) A scatter plot of modulus vs. hardness for a 1 mg/ml SSIP coating, with data points collected at different temperatures showing the temperature-dependent mechanical behavior. (b) Distribution curves of the modulus for SSIP coatings measured at temperatures ranging from 23°C to 170°C. (c) Hardness distribution of SSIP coatings across the same temperature range. (d) Surface maps of hardness (left) and modulus (right) at various temperatures for SSIP coatings, providing a visual representation of the spatial distribution of mechanical properties. (e) Comparative scatter plot and distributions of modulus for SSIP coatings at 140°C, highlighting the effect of concentration on the mechanical characteristics, with a control (uncoated substrate) for reference.

Mirroring the hydrophobicity results, the nanomechanical analysis indicates that SSIP films at a concentration of 1 mg/mL exhibit an increase in modulus and hardness upon exposure to increasing thermal treatment, reaching to a maximum at 140 °C (**Fig. 4**). At this juncture, the SSIP films exhibit an increase in modulus from approximately 135 GPa at 23 °C to 155 GPa at the 140 °C treatment with a noteworthy increase of approximately 15%. Similarly, the hardness progresses from nearly 5 GPa to a peak of 9 GPa within the same temperature range, signifying an increase of around 80%. This peak corresponds with the optimal coating condition identified through contact angle measurements, suggesting a unified optimal condition for enhancing both the hydrophobic and mechanical characteristics of the coatings (**Fig. 4**).

Conversely, at higher concentrations, this includes 5 mg/mL and 10 mg/mL, this trend attenuates, and a plateau in mechanical properties is observed, despite further thermal treatment. This plateau mirrors the convergence seen in hydrophobicity at high SSIP concentrations and elevated temperatures, highlighting a potential saturation point beyond which additional SSIP or heat fails to significantly reinforce the coating’s properties. The pronounced increase in both hardness and modulus at optimal conditions (1 mg/mL SSIP concentration at 140 °C) could imply a thermally induced rearrangement of the protein structure, leading to a denser, possibly more cross-linked network.^[28–30]^ This hypothesis aligns with the speculation of heat-induced denaturation and aggregation contributing to enhanced hydrophobic domains, as previously discussed. At higher SSIP concentrations, the observed lack of continued increase in mechanical properties could be attributed to similar phenomena that affect hydrophobicity. Overcrowding of proteins might lead to a less efficient packing structure or expose hydrophilic regions, thereby diminishing the nanomechanical integrity. The potential for increased porosity with thickness could also result in trapped solvent molecules, impacting the film’s mechanical response.

Furthermore, when we examined the SSIP films at the lower concentrations of 0.01 mg/mL and 0.1 mg/mL we noted the values closely resemble those of the uncoated silicon substrate, with the modulus being approximately 140-160 GPa and the hardness around 9-10 GPa. The mechanical properties of the uncoated silicon substrate can be considered a baseline, with a typical modulus of about 163 GPa and hardness in the range of 10 GPa (**Fig. 4**). This suggests that the SSIP layer at these concentrations contributes minimally to the overall mechanical response, with the observed values being predominantly representative of the substrate’s properties. The similarity in modulus and hardness between the low-concentration SSIP films and the silicon control could indicate that the SSIP layer thickness is insufficient to create a measurable mechanical distinction. The substrate effect is likely at play here, where the inherent properties of the silicon substrate govern the nanoindentation response due to the minimal thickness of the SSIP coating. The indentation size effect is also a consideration, as the indentations at these low concentrations may penetrate through the thin SSIP layer to the silicon substrate beneath, thereby influencing the results. If the indentation depth is similar to or greater than the thickness of the SSIP coating, the measured mechanical properties will be skewed towards those of the underlying silicon, overshadowing the true properties of the SSIP material. As the concentration increases to 1 mg/mL, there is a notable divergence from the substrate’s properties, with both modulus and hardness decrease, indicating that a more substantial SSIP layer contributes to the film’s mechanical behavior. At this optimal concentration, the SSIP coating likely forms a continuous and more uniform layer that can effectively manifest its inherent mechanical properties, minimizing the influence of the silicon substrate.

In summary, our nanomechanical data indicate an optimal SSIP concentration and thermal treatment that maximize the coating’s mechanical properties. This optimization is likely a result of balanced protein coverage and molecular reconfiguration, including heat-induced cross-linking or rearrangement and altered film density, which enhance the film’s structural integrity and mechanical characteristics.^[28–30]^ Comprehending these dynamics is essential for crafting SSIP coatings with mechanical properties tailored to specific applications, ensuring the SSIP layer effectively contributes without the substrate’s undue influence or the constraints of excessive bulk.

### 3.5 Variations in mechanical properties of color-differentiated SSIP film

Most of our experimental observations yielded a multicolored film upon casting SSIP. The SSIP film exhibited a colorful interference pattern, presumably indicative of varying film thickness. The colors range from red, corresponding to the thickest part of the film at approximately 700 nm, down to dark blue, representing the thinnest areas at around 400 nm (**Fig. 5**). These color changes are expected to be due to thin-film interference, where different film thicknesses alter the path of light waves, causing constructive or destructive interference that appears as a spectrum of colors to the observer.^[35,36]^ Given the heterogeneous mechanical landscape as depicted by the nanoindentation results described earlier, we hypothesis a correlation with the coloration pattern due to film thickness variations. Understanding the relationship between film thickness, mechanical properties, and visual appearance is essential for tailoring the fabrication process to achieve the desired film uniformity and performance characteristics in the future. ^[37]^ These variances in mechanical behavior are critical to understanding as they may impact the film’s performance in applications where uniformity in mechanical properties is important to understanding the intrinsic material behavior under varying film thicknesses, which could affect the packing density, degree of cross-linking, and presence of structural defects or inhomogeneities within the film.

**Figure 5.**
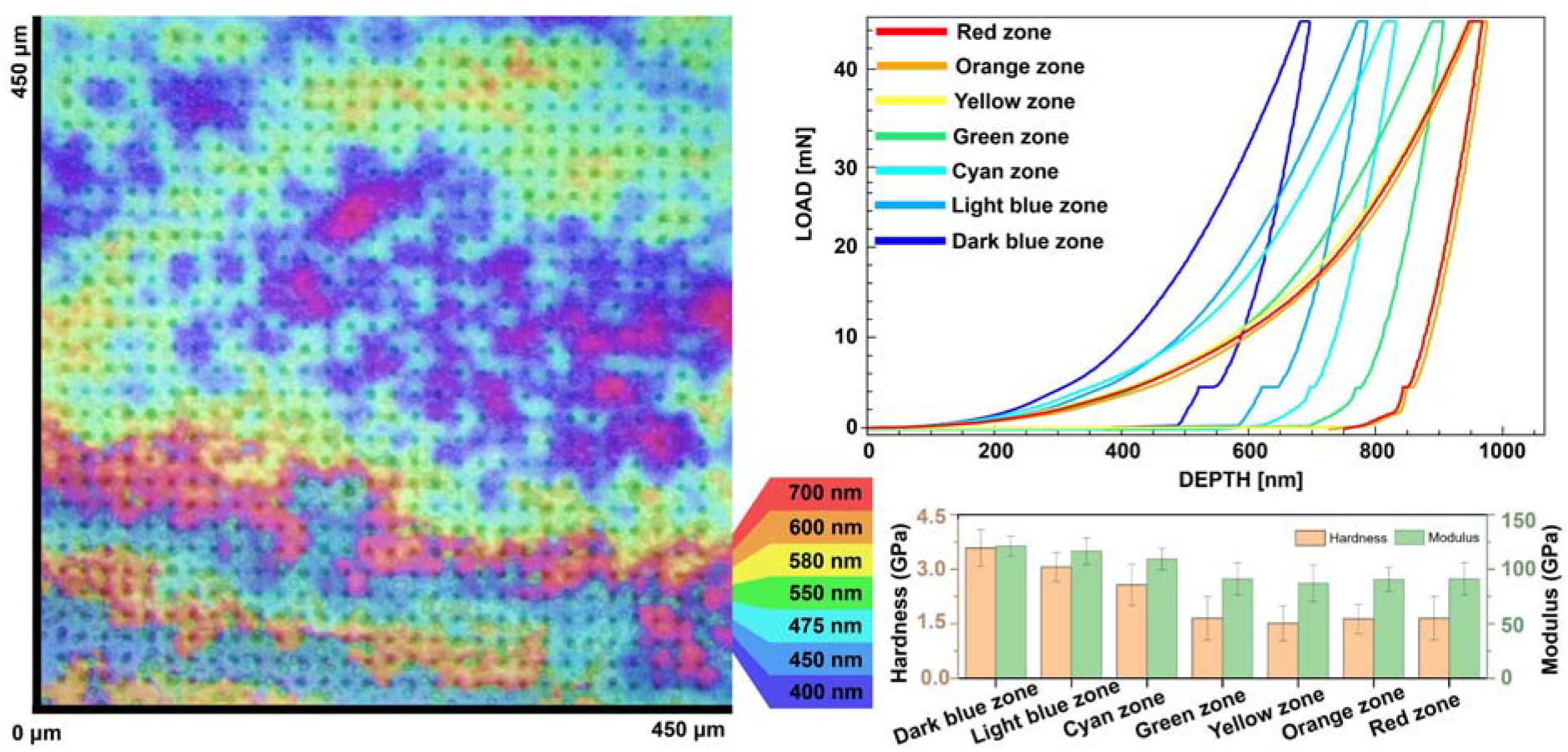
Representation of the mechanical properties of the film across different color zones, each corresponding to a certain thickness as denoted by the legend. Figure also shows load vs. depth curves for each color-coded zone, indicating material response to compressive force at varying depths as well as bar graphs representing the calculated hardness and modulus for each zone, reflecting the mechanical diversity within the material.

To this end, we carried out additional nanoindentation across these color-differentiated zones to identify quantitative insights into the mechanical properties of the SSIP film. The nanoindentation results illustrate the mechanical responses associated with thickness variations. The indentation curves suggest that the mechanical properties of the film are heterogeneously distributed with dark blue exhibiting higher resistance to indentation compared to the red areas. The load-depth curves, aligned with the film’s color map, indicate that the mechanical resistance of the film varies with the color zones, which are indicative of film thickness. The hardness values decrease as the film thickness increases. The dark blue zone shows higher hardness, near 4.5 GPa, while the red zone, which is thicker, has a lower hardness, close to 1.5 GPa (**Fig. 5**). This suggests a thickness-dependent decrease in hardness, with the thicker red zone being roughly 67% softer than the thinner dark blue zone. The modulus values demonstrate a similar pattern. The modulus is highest in the dark blue zone, approximately 150 GPa, and decreases in the red zone to about 50 GPa, indicating that the thicker regions have a modulus that is 67% lower than the thinner regions (**Fig. 5**). This inverse relationship between thickness and mechanical properties is intriguing, as it suggests that thinner films may have a denser, possibly more cross-linked structure, which often correlates with increased hardness and stiffness. The exact mechanisms driving these results would require further molecular-level analysis, potentially involving the evaluation of the film’s microstructure, degree of polymerization, and cross-linking density.

## 4. Conclusion

The exploration and development of advanced material coatings represents a pivotal advancement in the field of materials science and engineering, embodying a paradigm shift towards the integration of innovative technologies and sustainable methodologies. This transformative journey is underscored by a deliberate move away from the reliance on petrochemical-derived materials, steering towards the utilization of bio-based components that promise not only environmental sustainability but also a leap in material performance. Central to this innovation we propose the utilization of biosynthetically produced high-performance components. Leveraged through the advancements in recombinant DNA technology, SSIPs exemplify the fusion of biological inspiration with material innovation, offering a sustainable alternative to traditional coatings. The distinctive attributes of SSIPs, including their ease of processing, benign environmental footprint, and exceptional physicochemical properties, mark a significant stride towards the realization of green chemistry principles in material development. The comprehensive insights from this research illuminate the intricate dynamics that govern the properties of SSIP-based coatings. The interplay between material concentration, thermal treatment, and physical structure is highlighted, paving the way for tailored fabrication processes. These processes could optimize SSIP coatings for a wide array of applications that demand specific hydrophobic and mechanical characteristics, thereby broadening the scope of their applicability. We envision advancement of SSIP-based coatings represents a significant leap towards the combination of sustainability and high performance in material coatings. By harnessing the unique properties of bioengineered materials and fine-tuning the fabrication parameters, this study not only contributes to the field of materials science and engineering but also underscores the potential of sustainable innovations in addressing the current material challenges. The use of high performance biologically derived component has the promise of revolutionizing material coatings and holds a vast implication for various fields, including electronics, where the demand for high-performance, environmentally benign materials is ever-increasing.

## Credit authorship contribution statement

Anni Seisto: Investigation, Validation. Anna S. Borisova: Conceptualization, Validation. Robert Pylkkänen: Visualization. Pezhman Mohammadi: Conceptualization, Investigation, Validation, Supervision.

## Data availability

Data will be made available on request.

## Acknowledgements

This work was supported by the Academy of Finland project 348628, and internal funding from the VTT Technical Research Center of Finland, Ltd.

